# Competition Between Task Accuracy and Habitual Locomotor Behavior Underlies the Speed-Accuracy Trade-Off in Walking

**DOI:** 10.64898/2026.10.21.760535

**Authors:** Isaiah J. Lachica, Nicolas Schweighofer, James M. Finley

## Abstract

The speed-accuracy trade-off is a hallmark feature of motor behavior, yet its underlying causes in cyclic, whole-body movements such as walking remain to be understood. In discrete, point-to-point movements, reduced accuracy at higher speeds is typically attributed to signal-dependent motor noise and an inability to use online error correction. Walking, however, may be governed by additional objectives, such as minimizing metabolic energy expenditure. To distinguish between these mechanisms, we tested young adults in a precision-walking task requiring them to match target step lengths across a range of treadmill speeds. Accuracy declined systematically as walking speed increased, demonstrating a locomotor speed-accuracy trade-off. Contrary to prevailing theory, removing online visual feedback of foot position did not affect this decline, indicating little contribution from visually guided error correction. Moreover, step-length variability, a task-specific measure of signal-dependent motor noise, decreased rather than increased with speed. Instead, faster walking produced progressively larger biases toward participants’ habitual step lengths measured in the absence of targets. Because habitual step lengths are known to closely approximate energetically optimal values, these findings suggest energetic optimization, rather than motor noise or limitations associated with the use of online feedback, as the primary driver of the speed-accuracy trade-off during walking.

**New and Noteworthy:** Here we examine factors that contribute to the speed-accuracy trade-off during precision walking. We found that foot placement accuracy declined at faster walking speeds, and this decline was accompanied by a progressive increase in bias towards participants’ habitual step lengths. Because habitual stepping strategies approximate energetically optimal values, these findings suggest that the speed-accuracy trade-off during precision walking may emerge from competition between error minimization and energetic optimization.

## Introduction

When walking in complex environments, such as cluttered paths or natural terrain where missteps could result in falls, we naturally walk slower^1,2^ to improve foot placement accuracy^3,4^. Similarly, when walking speed is constrained, errors between desired and actual foot placement increase with walking speed^5^. This speed-accuracy trade-off is a fundamental principle of motor behavior^6–10^. In particular, studies of upper extremity tasks and saccadic eye movements have identified two primary factors thought to drive this trade-off. First, the ability to perform visually-guided online error corrections becomes increasingly limited as movement speed increases due to visuomotor processing delays^9–15^. Second, faster movements are also associated with greater signal-dependent motor noise^16,17^, observed as force variability increasing with force magnitude^7,18–21^. Because fast movements are generated with large forces, they produce greater movement variability and, ultimately, greater errors. While online error correction limits and signal-dependent motor noise produce a speed-accuracy trade-off in discrete movements, it remains unclear whether these factors contribute to the trade-off in precision walking.

Prior work on discrete lower-extremity movements suggests that online error correction limits may contribute to the greater foot placement errors when walking at faster speeds. Foot placement errors were greater in fast than slow steps during discrete stepping movements^22,23^. Visually guided corrections also occurred later and closer to targets in fast steps, reducing the time and space for online error corrections^23^. Signal-dependent motor noise during walking has been quantified by measuring changes in the variability of push-off force, joint torques, and step length variability as walking speed increases. While push-off force^24,25^ and push-off force variability^25,26^ increase monotonically with walking speed, step length variability has been found to either decrease with speed^25^ or remain unchanged by speed^27^. While associations between variability and speed have been demonstrated during steady-state walking, it remains unclear whether they would extend to precision walking and contribute to a speed-accuracy trade-off.

Because walking is a continuous, cyclical behavior, its dynamics may influence the speed-accuracy trade-off in ways distinct from discrete movements. In particular, step length is naturally linked to walking speed, with people taking longer steps as they walk faster^24,28^. This habitual relationship between step length and speed typically minimizes the energy needed to move a given distance^29–33^. Thus, deviations from habitual step lengths would increase energy cost, and this cost would increase with walking speed^34,35^. The energy cost of accurately hitting a target step length can thus be thought of as a function of the difference in energy cost between the target and the habitual step length. Prioritizing energy minimization would increase step length error as the difference between target and habitual step lengths grows. As such, the speed-accuracy trade-off could arise from biasing step lengths toward their habitual values to minimize energy costs, particularly at faster speeds when energy cost penalties are higher.

Our objective was to determine the factors that contribute to the speed-accuracy trade-off during precision walking. Fifteen young adults completed a treadmill-based precision walking task, matching their step lengths to target step lengths displayed on a screen while walking at increasing fixed treadmill speeds. We hypothesized that the speed-accuracy trade-off extends to our precision walking task, with step length error increasing with walking speed. We also hypothesized that speed-dependent limits on visually guided online error corrections partially contribute to the trade-off during precision walking. To test this hypothesis, participants completed the task with full and endpoint feedback conditions. In the former, participants received visual information about foot position throughout the swing phase, whereas in the latter it was available only at heel strike, preventing visually-guided online error corrections across all walking speeds. We predicted that error would increase less as walking speed increased in the endpoint feedback condition. We also hypothesized that signal-dependent motor noise does not contribute to the trade-off during precision walking. To test this hypothesis, we quantified changes in step length variability, a task-specific indirect measure of signal-dependent motor noise, as walking speed increased. Consistent with prior observations during steady-state walking, we predicted that step length variability would decrease as walking speed increased. Finally, we hypothesized that a speed-dependent shift toward a habitual stepping strategy to minimize energy cost also contributes to the speed-accuracy trade-off during precision walking. To test this hypothesis, we modeled step length errors as a function of target and habitual step lengths. We predicted that step lengths would be increasingly biased toward habitual step lengths as walking speed increased.

## Methods

### Participants

We recruited 15 young adults (27.3 ± 3.5 years, 9F/6M) with no musculoskeletal or neurological injuries that could have affected their ability to walk and complete the task. Study procedures were reviewed and approved by the University of Southern California’s Institutional Review Board. Participants provided written informed consent before participating and received monetary compensation for their time. All aspects of the study conformed to the principles described in the Declaration of Helsinki.

### Precision Walking Task

Participants completed a treadmill-based precision walking task in which they matched their step lengths to targets displayed on a screen while walking at five fixed-speed conditions corresponding to 60%, 80%, 100%, 120%, and 140% of their self-selected walking speed. The participants were instructed to hit each target accurately by placing a white dot, representing foot position calculated as the anteroposterior distance between the feet, in the center of blue rectangles, representing the target step lengths (Fig. 1A-B). We identified each participant’s self-selected walking speed using a custom staircase algorithm. First, participants walked on the treadmill at 75% of their overground walking speed, determined as the average speed from three 10-meter walk tests. We then increased or decreased treadmill speed in 0.05 m/s increments and computed self-selected walking speed as the average of the four speeds participants indicated as comfortable.

**Figure 1.**
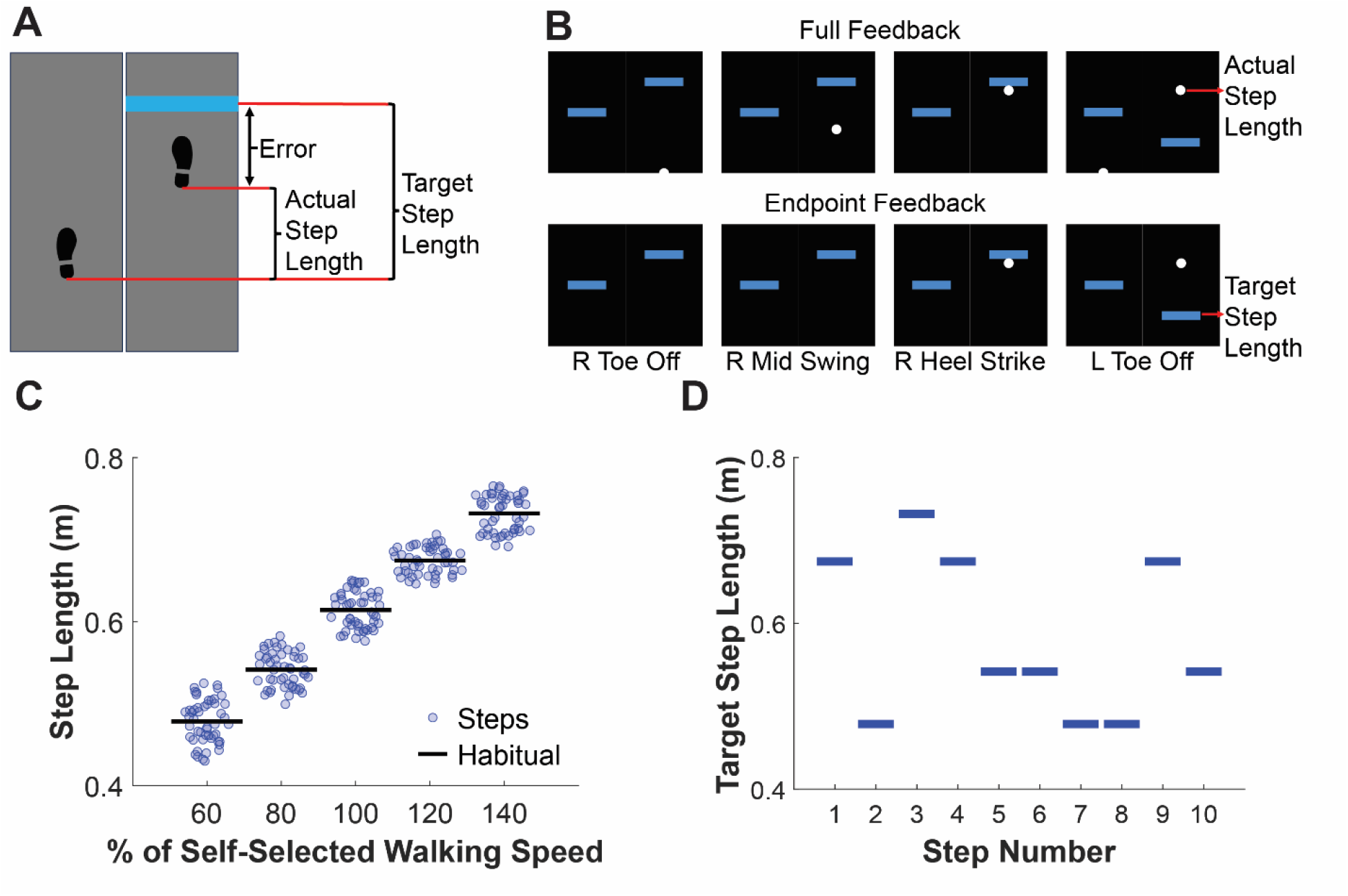
Experimental design. (A) Top-down view of the precision walking task on the treadmill showing the difference between actual and target step lengths. (B) Visual feedback during four phases of a step. Foot position, calculated as the anteroposterior distance between the feet, was shown as a white dot on the screen, while target step lengths were shown as blue rectangles. Actual step lengths corresponded to the distance between the feet at heel strike. Visual information about foot position was provided throughout the swing phase in the full feedback condition (upper row) and only at heel strike in the endpoint feedback condition (lower row). (C) Data from a representative participant illustrating their habitual step lengths which was calculated as the average of 50 steady-state steps at each fixed-speed condition. (D) Target step lengths for the first 10 steps which was randomly sampled from the set of habitual step lengths for the same representative participant in C.

Before the precision walking task, participants completed a separate steady-state walking task at each of the five fixed-speed conditions. We calculated their habitual step length at each fixed speed condition as the average of 50 steady-state steps, resulting in five habitual step lengths (Fig. 1C). Target step lengths during the precision walking task were then drawn from this set of habitual step lengths. Specifically, we replicated each habitual step length 50 times to generate 250 target step lengths, then randomly shuffled them once to generate a single sequence used in every trial (Fig. 1D).

To test whether speed-dependent limits on the ability to perform visually-guided online error corrections due to visuomotor processing delays contribute to the trade-off during precision walking, the participants completed the task in two feedback conditions (Fig. 1B). In the full feedback condition, the participants were provided with visual information about their foot position starting from toe-off, throughout the entire swing phase, and ending at heel strike. In the endpoint feedback condition, participants received visual information about their foot position only at heel strike, limiting their ability to perform visually-guided online error corrections.

Participants completed four practice trials, each with 100 targets. During the practice trials, participants familiarized themselves with the on-screen visual feedback and practiced using it to adjust their stepping strategy and match target step lengths. In the first practice trial, participants walked at 100% of their self-selected speed and were presented with target step lengths corresponding to their habitual step length at that speed. This first practice trial was designed to familiarize participants with the various components of the visual feedback on the screen. In the second and third practice trials, participants practiced with the most challenging target step lengths at the slowest and fastest speeds. In the second practice trial, participants walked at the slowest speed and encountered the longest target step lengths. In the third practice trial, participants walked at the fastest speed and encountered the shortest target step lengths. In the final practice trial, participants walked at 100% of their self-selected walking speed while matching target step lengths generated by combining 20 replicates of each habitual step length, for a total of 100 randomly shuffled targets.

Participants then completed the test trials, consisting of two sets of five fixed-speed trials, with each set corresponding to one feedback condition and each fixed-speed trial corresponding to one of the five speeds, for a total of 10 test trials. The feedback condition sets were counterbalanced across participants, and the fixed-speed trials were randomized within each set to account for potential order effects. Before each test set, participants completed the four practice trials described above, modified to match the feedback condition they would experience.

### Data Collection and Processing

We used an eight-camera motion capture system (Qualisys AB, Gothenburg, Sweden) sampling at 100Hz to collect three-dimensional kinematic data from eight passive reflective markers to track lower extremity movements. The passive reflective markers were placed on the bilateral greater trochanters, lateral malleoli, distal phalanges of the first toes, and heels using an eight-camera motion capture system. We also collected kinetic data using a split-belt instrumented treadmill (Bertec Corporation, Columbus, OH, USA), sampling at 1000Hz for gait event detection. We then streamed kinematic and kinetic data in real time from Qualisys Track Manager to MATLAB (MathWorks, Natick, MA, USA) to run our custom visual feedback code.

We defined heel strike as the time when the vertical component of ground reaction force was greater than or equal to 80N. We quantified step length as the anteroposterior distance between the bilateral heel markers at each heel strike. We then calculated step length error as the average absolute difference between the actual and target step length across all steps in a trial. We also used step length variability as a task-based indirect measure of signal-dependent motor noise by calculating the standard deviation of all step lengths in a trial. We calculated step length error and step length variability in each fixed-speed and feedback condition.

### Statistical Analysis

All statistical analyses were performed in R (R Project for Statistical Computing). We used the *lme4* package to fit linear mixed-effects models to our data. We also used the *robustlmm* package to fit robust linear mixed-effects models to address violations of normality and homoscedasticity in the residuals as well as to account for outliers and influential points as determined by Cook’s distance.

#### Characterizing the speed-accuracy trade-off during precision walking

To test for a speed-accuracy trade-off in our precision walking task and determine whether the speed-dependent limitations on visually-guided online error corrections contributed to the trade-off, we fit a robust linear mixed-effects model with step length error as the dependent variable and speed, feedback condition, and their interaction as predictors. We included a random intercept for each participant to account for repeated measures and inter-participant variability. To test whether signal-dependent motor noise contributed to the trade-off, we fit a robust linear mixed-effects model with step length variability as the response and speed, feedback condition, and their interaction as predictors. We included a random intercept for each participant in the step length variability model to account for repeated measures and inter-participant variability. A log-likelihood ratio test also indicated that adding a random slope for speed improved fit to the step length variability model. To assess the statistical significance of the regression parameters, we computed 95% confidence intervals via bootstrapping and checked whether each interval included zero. We first generated 1000 bootstrap datasets by resampling participants with replacement. To replicate the distribution of the original data, each resampled dataset contained 15 participants with data from all fixed speed conditions and from both feedback conditions. We then fit the models to each dataset, extracted the coefficient estimates for each predictor, and calculated the 95% confidence intervals using the bias-corrected and accelerated (BCa) percentile method in the *coxed* package.

#### Modeling the influence of gait-specific factors on precision walking performance

To test the influence of energy minimization on the speed-accuracy trade-off during precision walking, we fit linear mixed-effects models using data from the full-feedback condition. We modeled actual step lengths (SL^A^) as a function of target step length (SL^T^), which reflects the objective of maximizing stepping accuracy; habitual step length specific to the current speed condition (SL^H^), which represents what participants would do if their objective was to minimize energy cost; and their interaction. We included a random intercept for each participant and random slopes for SL^T^, SL^H^, and their interaction to account for individual differences in how these predictors affected SL^A^. We modeled random intercepts and slopes independently. We fit reduced versions of the full model to assess each predictor’s relative importance (Table 1). We considered the most influential predictor to be the one whose omission yielded the greatest change in both BIC and marginal R^2^ relative to the full model. In the models, the subscript *n* denotes the current step, while *s* denotes the current fixed-speed condition. Predictors in the full model were considered significant if their p-values were less than the alpha value set at 0.05.

**Table 1.** Modeling actual step lengths (SL^A^) in the full feedback condition as a function of the habitual step length specific to the current speed condition (SL^H^), target step length (SL^T^), and their interaction. Reduced models, where each predictor was systematically omitted from the full model, were fit to determine the relative importance of each predictor to model fit.

| Model name | Model structure |
|---|---|
| Habitual step lengths | SL<sup>A</sup><sub>n</sub> ~ SL<sup>H</sup><sub>s</sub> |
| Target step lengths | SL<sup>A</sup><sub>n</sub>~ SL<sup>T</sup><sub>n</sub> |
| Habitual and target lengths | SL<sup>A</sup><sub>n</sub>~SL<sup>T</sup><sub>n</sub> + SL<sup>H</sup><sub>s</sub> |
| Full model with interaction between habitual and target lengths | SL<sup>A</sup><sub>n</sub>~ SL<sup>T</sup><sub>n</sub> + SL<sup>H</sup><sub>s</sub> + SL<sup>T</sup><sub>n</sub>x SL<sup>H</sup><sub>s</sub> |

## Results

### Online error correction limits and the speed-accuracy trade-off

We found a main effect of walking speed, as step length errors increased with walking speed (*β* = 0.027 m, 95% CI = [0.020 m, 0.036 m]; Fig. 2). We also found a main effect of feedback condition, with greater step length errors in the endpoint feedback condition than the full feedback condition (*β* = 0.015 m, 95% CI = [0.004 m, 0.031 m]). However, we did not find an interaction between walking speed and feedback condition, as the increase in step length error with walking speed was similar in both feedback conditions (*β* = -0.006 m, 95% CI = [-0.016 m, 0.005 m]). While these results show that the speed-accuracy trade-off extends to our precision walking task, they also suggest that online error correction limits do not contribute to the trade-off.

**Figure 2.**
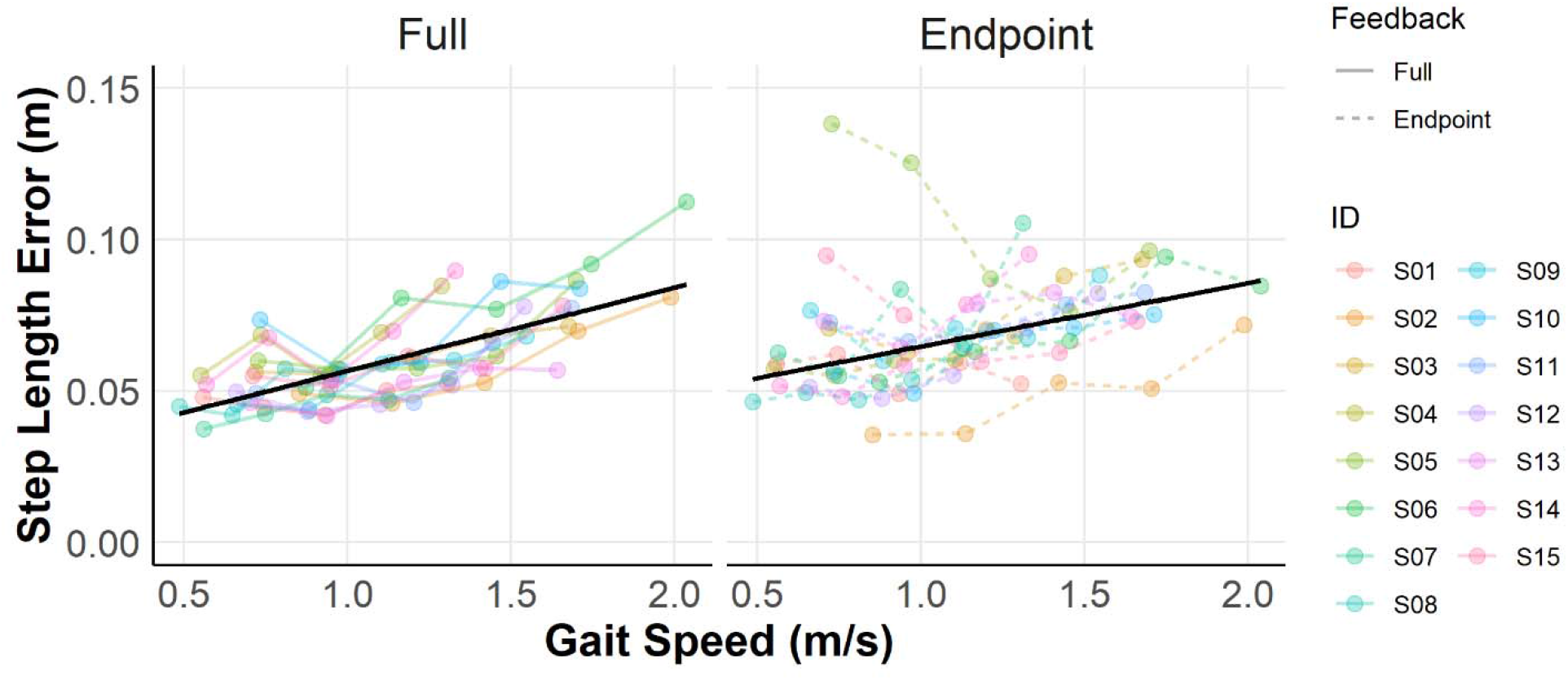
Effect of gait speed on step length error in the full and endpoint feedback conditions. Solid black lines correspond to fixed-effect model fits (Marginal R^2^ = 0.42). Errors increased similarly with gait speed in both the full-feedback and endpoint-feedback conditions.

### Signal-dependent motor noise and the speed-accuracy trade-off

We found a main effect of walking speed on step length variability (Fig. 3). However, this effect was in the opposite direction of what would be expected if step lengths were influenced by signal-dependent motor noise. Specifically, step lengths became less variable as gait speed increased (*β* = -0.010 m, 95% CI = [-0.018 m, -0.001 m]). We also found a main effect of feedback condition, with step length variability greater in the endpoint feedback than in the full feedback condition (*β* = 0.022 m, 95% CI = [0.015 m, 0.034 m]). However, there was no interaction between walking speed and feedback condition, indicating that the decrease in step length variability as walking speed increased was comparable between the full feedback and endpoint feedback conditions (*β* = -0.004 m, 95% CI = [-0.013 m, 0.003 m]). Thus, the results indicate that signal-dependent motor noise does not contribute to the trade-off during precision walking.

**Figure 3.**
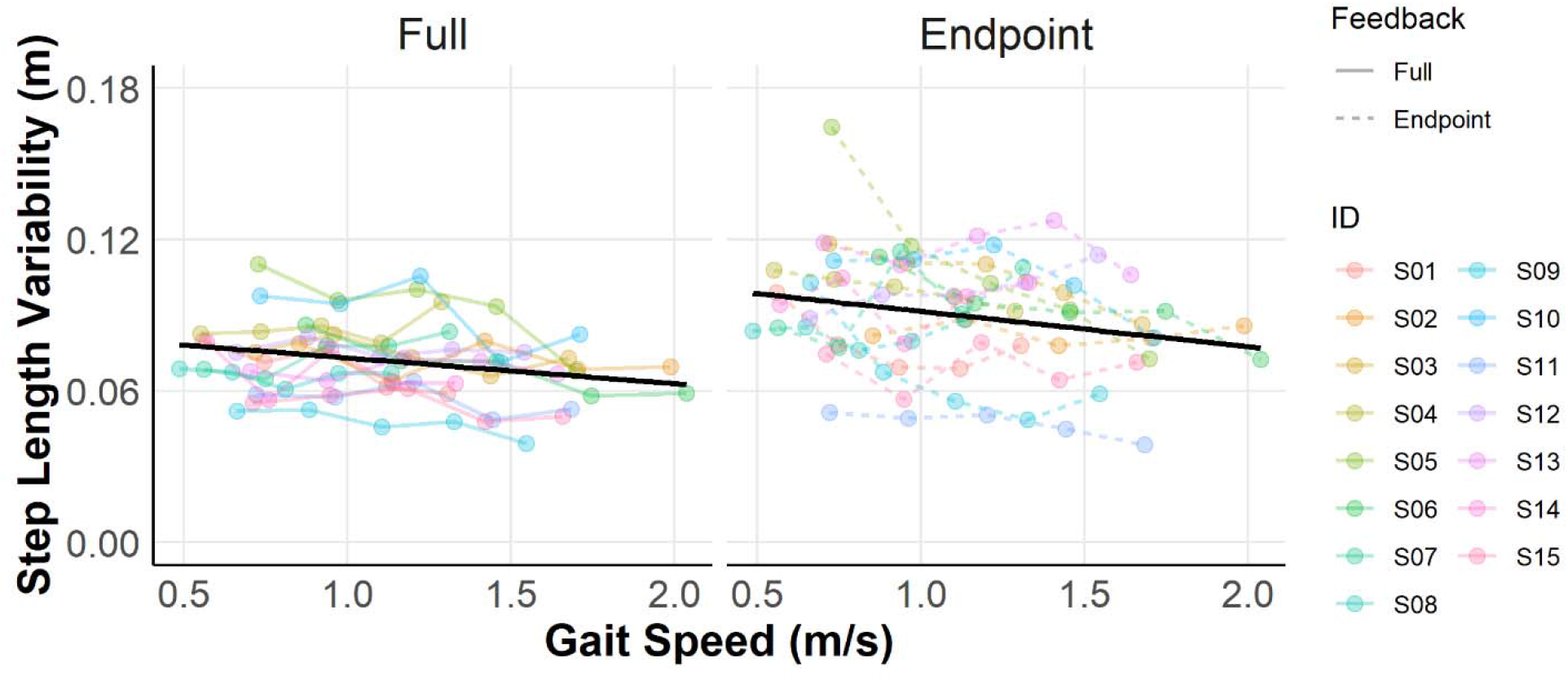
Effect of gait speed on step length variability in the full-feedback and endpoint-feedback conditions. Solid black lines represent fixed-effect model fits (Marginal R^2^ = 0.50). Step length variability decreased as gait speed increased in both conditions, though it was lower in the full feedback condition than in the endpoint feedback condition.

### Energy cost minimization and the speed-accuracy trade-off

We found that the full model, which included an interaction between target and habitual step length, fit the data best, as indicated using BIC and log-likelihood ratio tests when compared with each reduced model (Table 2). Among the predictors, SL^T^ appeared to be the most important to the goodness of fit of the full model, as its omission resulted in the greatest change in BIC (ΔBIC = 9617) and marginal R^2^ (ΔMarginal R^2^ = 0.35).

**Table 2.** Model comparison results. * Indicates statistically significant log-likelihood ratio tests comparing each reduced model to the full model. BIC = Bayesian Information Criterion; Marginal R² = variance explained by fixed effects alone.

| **Model name** | **BIC** | **Log-likelihood** | **Marginal R<sup>2</sup>** |
|---|---|---|---|
| Habitual step lengths* | -44806 | 22427 | 0.04 |
| Target step lengths* | -52383 | 26216 | 0.21 |
| Habitual and target step lengths* | -54007 | 27038 | 0.36 |
| Full model with interaction between habitual and target lengths | -54423 | 27256 | 0.39 |

In the full model, SL^T^ (*β* = 1.07, p < 0.001), SL^H^ (*β* = 0.75, p < 0.001), and their interaction (*β* = -0.98, p < 0.001) were all significant predictors of SL^A^ (Fig. 4A). The fixed effects of the full model alone accounted for 38% of the variance in the data (Marginal R^2^ = 0.38) while the random effects accounted for an additional 34% based on individual-specific variability particularly for SL^H^ and its interaction with SL^T^ (Conditional R^2^ = 0.72, Fig. 4A & 4B). Taken together, these model results indicate that participants weighted SL^T^ more heavily than SL^H^ when SL^H^ was smaller. However, the influence of SL^T^ decreased as SL^H^ increased, which occurred at faster walking speeds. This relationship is best illustrated in the adjusted prediction plots in Fig. 4C, which illustrate the relationship between SL^T^ and SL^A^ while holding SL^H^ constant. Here, the selected SL^H^ values were based on the average SL^H^ across all participants at each of the five fixed-speed conditions. The slope between SL^T^ and SL^A^ was steeper and closer to the unity line (dotted black line) at smaller SL^H^ values (SL^H^ = 0.47 m), indicating more accurate step lengths. As SL^H^ increased, the slope between SL^T^ and SL^A^ became progressively shallower and moved away from the unity line, which contributed to greater step length errors (Fig. 2).

## Discussion

We used a treadmill-based precision walking task to determine the factors contributing to the speed-accuracy trade-off during precision walking. Consistent with recent studies^5,22,23^, we observed a speed-accuracy trade-off in our task: as walking speed increased, step length errors also increased. Although removing online visual information increased overall error, errors increased with speed similarly with and without online feedback about step lengths. These results suggest that while visual information about the effector is critical for accuracy, the increase in error with speed cannot be explained by speed-dependent limits on the ability to perform visually-guided online error corrections due to visuomotor processing delays. Additionally, step length variability decreased as walking speed increased in both feedback conditions, suggesting that signal-dependent motor noise may not have contributed to the trade-off observed in our task. Instead, we found that participants produced step lengths increasingly biased toward habitual step lengths as walking speed increased. Together, these results suggest that a speed-dependent shift away from matching targets and toward a habitual stepping strategy, seemingly to prioritize energy cost minimization over stepping accuracy, primarily contributed to the trade-off observed.

**Figure 4.**
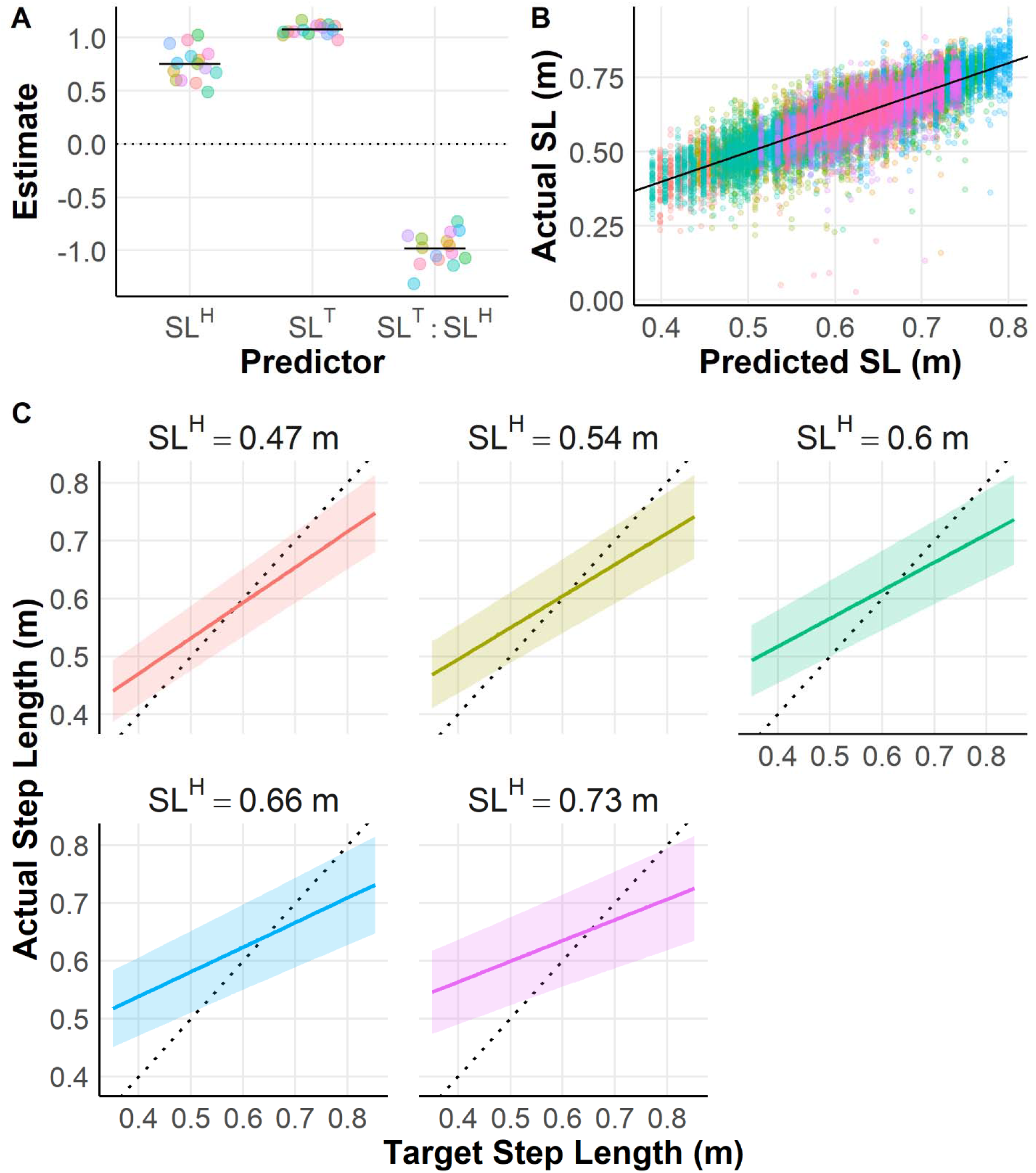
Mixed-effects model results for SL^A^ as a function of SL^T^, SL^H^, and their interaction. (A) Coefficient estimates for each predictor. Horizontal lines indicate the fixed-effect estimates (SL^H^ β = 0.75, SLT β = 1.07, SL^H^:SL^T^ β = −0.98; all p < .001), while each point represents the individual-specific coefficient estimate (fixed- and random-effects). (B) Actual vs. predicted SL^A^ from the full model, with each color representing data from an individual participant (Marginal R^2^ = 0.39, Conditional R^2^ = 0.72). The black line represents the unity line where predicted step lengths match the actual step lengths. (C) Adjusted prediction plots of SL^A^ as a function of SL^T^ at five levels of SL^H^ corresponding to the average SL^H^ across participants at each fixed speed condition. Shaded regions represent 95% confidence intervals. The dotted black line represents the unity line where SL^A^ matches SL^T^. Predicted SL^A^ was closest to the unity line at the smallest SL^H^ value (SL^H^ = 0.47 m) and moved progressively away from it as SL^H^ increased.

One of the factors proposed to primarily contribute to the speed-accuracy trade-off is the speed-dependent limit on the ability to perform visually-guided online error corrections due to visuomotor processing delays^9–15^. However, we found no evidence that these online error correction limits contributed to the trade-off during precision walking. On average, step length errors were greater in the endpoint feedback condition than in the full feedback condition. This finding is consistent with prior work showing that the accuracy of aimed movements decreases when visual information about the effector is decreased^12,14,36,37^ or removed completely^10,37,38^. However, we found that step length error increased with walking speed similarly in the endpoint feedback and full feedback conditions. Although this contrasts with prior work on discrete lower-extremity movements, our findings align with the literature on the role of gaze during walking. We often do not rely on online visual information for foot placement. Instead, we perform steps primarily in a feedforward, ballistic manner, with minimal need for online visual feedback. In particular, during precision walking in rough or complex terrain, visual information about foot placement targets is sampled approximately two steps in advance^3,4,39–42^. As such, online error correction limits may not contribute to the speed-accuracy trade-off in precision walking.

Another factor hypothesized to primarily contribute to the speed-accuracy trade-off is signal-dependent motor noise^7,17^. Signal-dependent motor noise predicts that movement variability will increase with movement speed, ultimately resulting in greater errors. However, we found the opposite trend in our task, as step length variability decreased as walking speed increased. Although this trend is inconsistent with signal-dependent motor noise, it matches what is known in steady-state walking, where step length variability either decreases^25^ or remains unchanged^27^ as walking speed increases. Additionally, there is an energy-optimal step length for a given speed^29–33^ that the participants would need to deviate from to accurately hit the target step lengths in our task, resulting in more variable step lengths. As the energy cost of deviating from energy-optimal step lengths also increases with gait speed^34,35^, participants may have prioritized minimizing energy cost over stepping accuracy particularly at faster speeds, resulting in the more consistent step lengths observed.

While most prior work on this trade-off has used discrete aiming movements, walking is unique as it is a continuous, cyclical movement that exhibits speed-dependent factors that may influence the trade-off. To investigate these factors, we modeled actual step lengths as a weighted linear combination of the habitual step length at a given speed (representing energy cost minimization), the target step length (reflecting task demands), and their interaction. In general, participants weighted the target step length more heavily than their habitual step length when habitual step lengths were short. However, the influence of target step length decreased as habitual step length increased, corresponding to faster speeds. These model results suggest that participants prioritized meeting task objectives, such as maximizing stepping accuracy, over minimizing energy cost at slower speeds. However, prioritization shifted away from maximizing stepping accuracy as walking speed increased, potentially because the energy cost of deviating from energy-optimal step lengths increases at faster walking speeds ^34,35^. Overall, our results suggest that speed-dependent competition between adhering to task demands and minimizing energy cost may underlie the speed-accuracy trade-off during precision walking.

### Limitations

We indirectly evaluated the influence of signal-dependent motor noise on the speed-accuracy trade-off by measuring movement endpoint variability during gait using step length variability. One way to directly measure signal-dependent motor noise is to quantify variability in the forces generated during the movements of interest. In walking, future studies can examine variability in peak ground reaction force or in lower-extremity muscle activity, as measured by electromyography, during the swing phase. Similarly, we indirectly inferred the role of minimizing energy cost in the trade-off by modeling step lengths as a combination of the target and habitual step lengths at a given speed, referring to the well-established relationship between habitual step lengths and energy cost in the literature. Future work could directly measure energy cost by quantifying joint-level work, cost of transport, or both during the task. Lastly, other factors, such as balance control, may influence foot placement and the speed-accuracy relationship observed here. For example, deviations in both mediolateral and anteroposterior foot placement during unperturbed walking can be modeled as a function of the position and velocity of the body’s center of mass ^43,44^. If this control strategy remains in place during precision walking, it may account for additional variance in step length error not captured by the models evaluated here.

## Conclusion

Our findings suggest that neither speed-dependent limits on visually-guided online error corrections nor signal-dependent motor noise alone explains the speed-accuracy trade-off during precision walking. Instead, this trade-off appears to be driven by a speed-dependent competition between two objectives: minimizing energy cost and maximizing task performance. This influence of energy cost on the speed-accuracy trade-off may be unique to walking and may explain why previously established factors known to drive the trade-off in other movements may not contribute significantly to the trade-off during precision walking. These results highlight the importance of energy cost minimization as an objective function that explains multiple features of human walking beyond steady-state walking.

## Data Availability

The processed data and the code used to generate the figures and perform the statistical analyses are available on Zenodo^45^.

## CRediT Authorship Contribution Statement

**Isaiah J Lachica:** Conceptualization, Methodology, Software, Formal analysis, Investigation, Data curation, Visualization, Writing - original draft, Writing - review & editing. **Nicolas Schweighofer:** Writing - review & editing, Methodology. **James M Finley:** Conceptualization, Methodology, Resources, Writing - original draft, Writing - review & editing, Supervision, Project administration, Funding acquisition.

